# CAZyme domain architectures suggest fine-scale functional differentiation among anaerobic fungi and bacteria during lignocellulose conversion to volatile fatty acids

**DOI:** 10.64898/2026.02.09.704939

**Authors:** Christopher E. Lawson, Joel P. Howard, Thomas S. Lankiewicz, James B. Brown, Steven W. Singer, Héctor García Martín, Michelle A. O’Malley

## Abstract

Anaerobic fermentation with microbial communities (microbiomes) is an emerging platform for conversion of lignocellulosic biomass to biofuels and bioproducts. The process relies on diverse anaerobic microbes that interact to deconstruct and convert lignocellulosic biomass into a range of products, such as volatile fatty acids (VFAs), which can be achieved by arresting methanogenesis during fermentation. However, defining the distinct functional roles played by various fungi and bacteria during anaerobic biodegradation remains poorly understood. Here, we performed parallel enrichment experiments from cow faeces, goat faeces, and anaerobic digester sludge, selecting for fungal or bacterial dominated communities that convert sorghum biomass into VFAs. Subsequently we reconstructed metabolic networks across these enrichments based on recovered bacterial metagenome-assembled genomes (MAGs) and fungal isolate genomes and profiled their metabolic activity using metatranscriptomics to identify potential functional niches. Our findings implicate diverse bacteria affiliated with the Bacteroidales and Lachnospiraceae in the direct conversion of lignocellulosic biomass to propionate and butyrate, respectively, whereas *Neocallimastix*-dominated fungal enrichments converted lignocellulose to lactate, acetate and formate. Analysis of carbohydrate-active enzymes (CAZymes) revealed fine-scale differences between microbes that expressed unique multi-functional enzymes linking two or more CAZymes together with distinct carbohydrate binding motifs, implicating lignocellulose structure as a key driver of selection and niche differentiation. Most of these multi-functional enzymes localized complementary degradation functions together, likely conferring synergistic degradation effects within and between microbiome members. We anticipate that these findings will help inform efforts to develop synthetic microbiomes with tailored functionality for low-cost conversion of lignocellulosic biomass to fuels and bio-based chemicals.

## Introduction

Biomanufacturing of chemicals from renewable lignocellulosic biomass has the potential to offset society’s reliance on fossil fuels. One biomanufacturing strategy involves the use of anaerobic microbial communities (microbiomes) to deconstruct and convert lignocellulosic biomass into biogas (methane and carbon dioxide) via anaerobic digestion [1]. While biogas can be upgraded to produce bioenergy (i.e., renewable natural gas or electricity), its market value is low, often rendering anaerobic digestion not economically viable. Therefore, higher value chemicals, such as volatile fatty acids (VFAs; $1-5/kg selling price) represent an emerging target product for anaerobic digestion that can be achieved by arresting methanogenesis during fermentation. These products can be chemically upgraded to liquid biofuels [2] or used directly in consumer products, antimicrobials or industrial materials [3].

Several studies have demonstrated VFA production via arrested methanogenesis from different organic feedstocks [4]. Conversion of lignocellulosic biomass has largely been achieved by anaerobic fermentations using rumen contents as inocula, resulting in acetate, propionate, and butyrate (C2-C4 VFAs) as the main fermentation products in the absence of an external electron donor [5–7]. During fermentation, the microbiomes responsible for lignocellulose deconstruction and conversion have been reported as containing diverse ruminal bacteria [6, 7], with no studies reporting the presence of biomass degrading ruminal fungi, indicating that they are outcompeted by bacteria. While the overall metabolic processes mediated by ruminal bacteria during lignocellulose degradation to VFAs are generally understood, the specific metabolic networks and interactions driving *in situ* microbiome metabolism remain largely unresolved. Moreover, the contribution and specific metabolic activities expressed by ruminal fungi during lignocellulose deconstruction are only starting to emerge [8].

Here, we enriched five parallel anaerobic fungal and bacterial consortia derived from the cow and goat rumen and an anaerobic digester on sorghum substrate in chemically defined media selecting for VFA production. We quantified metabolite and gas production across the enrichments to examine functional differences between bacterial-dominated consortia and fungal-dominated consortia, as well as inoculum source. Subsequently we reconstructed metabolic networks across the enrichments based on recovered bacterial metagenome-assembled genomes (MAGs) and fungal isolate genomes and profiled their gene expression using metatranscriptomics. Our results identify specific organisms, carbohydrate-active enzymes (CAZymes), and metabolic pathways active in the deconstruction and conversion of sorghum biomass to VFAs. These results were used to infer key metabolic interactions responsible for producing specific VFA end products and activities that compete with VFA production. Together, these findings offer molecular-level insights on anaerobic microbiome function and will inform the rational design of microbiomes for improved lignocellulose conversion to valuable chemicals.

## Methods

### Enrichment and cultivation of anaerobic bacterial and fungal cultures on sorghum biomass

Anaerobic consortia enriched for biomass-degrading bacteria and biomass-degrading fungi (herein termed “enrichments”) were cultivated in anaerobic Hungate tubes on 1% (w/w) sorghum (2 mm mill size; Idaho National Laboratory) as a sole substrate in 10 ml of chemically defined M2 media [9] with a ∼85% N_2_, 12% CO_2_, and 3% H_2_ headspace and resazurin (0.1% w/v) added as a visual redox indicator. Cultures were initially inoculated inside an anaerobic chamber (AS-580, Anaerobe Systems) with 1 mL of either cow (5 g wet feces in 25 mL M2 media) or goat (3 pellets ∼2.5 g in 25 mL M2) fecal solutions, or with 0.5 mL of sludge from an anaerobic digester to a final volume of 10 mL M2 media in the presence of either 5 mM chloramphenicol (CH, selecting for fungi) or 5 mM 2-bromoethanesulfonate (BES, selecting for bacteria) and passaged every 3-4 days based on 10% (v/v) transfers. Enrichment cultures were maintained at a pH of 7-7.5 and a temperature of 39°C. This resulted in 3 bacterial enrichments from the cow, goat, and digester inocula (cow+BES, goat+BES, digester+BES), and 2 fungal enrichments from the cow and goat inocula (cow+CH, goat+CH). Cow and digester enrichments were passaged for 10 generations (∼40 days), while goat enrichments were passaged for 5 generations (∼20 days). Analytical measurements were made on biological triplicates from each of the 5 enrichments cultures daily over a 4-day period. DNA/RNA samples were collected on Day 3 for metagenomic and metatranscriptomic analysis (see methods below).

### Measurement of fermentation products and total pressure accumulation

Total pressure accumulation in the headspace of Hungate tubes was monitored daily using a pressure transducer method as described previously [10]. Quantification of liquid fermentation products, including formate, acetate, propionate, butyrate, valerate, succinate, and lactate, was performed on a 1260 Infinity high-performance liquid chromatography (HPLC) system equipped with an Aminex HPX-87H column (part no. 1250140, Bio-Rad) with an inline 0.22 μm filter (Part No. 50671551, Agilent) followed by a Micro-Guard Cation H guard column (Part No. 1250129, Bio-Rad, Hercules, CA, USA) before the analytical column. Samples were prepared by acidification to a concentration of 5 mM H_2_SO_4_, followed by incubation at room temperature for 5 min, centrifugation at 21,000 x g for 5 min, and filtration through a 0.22 µm polyethersulfone (PES) membrane filter into HPLC vials with 300 µl polypropylene inserts. Metabolites were separated using a mobile phase of 5 mM sulfuric acid at 50°C with a flow rate of 0.6 ml/min, a run length of 30 min, and an injection volume of 20 μl. Signals were measured using a variable wavelength detector (λ = 210 nm) and standard curves for each compound were made at 5 concentrations bracketing the range expected in the samples.

Quantification of hydrogen (H_2_) and methane (CH_4_) gas in the headspace of Hungate tubes were analyzed on a Thermo Fisher Scientific TRACE 1300 gas chromatograph (GC) equipped with a TracePLOT™ TG-BOND Msieve 5 A (Part No. 26003-6100, Thermo Fisher Scientific) and an Instant Connect Pulsed Discharge Detector (PDD) (Part No. 19070014, Thermo Fisher Scientific). 100 µL of headspace gas was collected and subsequently purged three times in a 100 µL air-tight syringe and needle. Then, 20 µL of headspace gas was collected and injected directly into the GC. The oven temperature for each run was 30 °C and the PDD temperature was 150 °C. High-purity helium (Part No. HE 5.0UH-55, Praxair, Danbury, CT, USA) was further purified with a heated helium purifier (Part No. HP2, VICI) and used as the carrier gas with a flow rate of 0.5 mL/min. The same flushing and analysis procedures were followed for methane and hydrogen standards including 500 ppm H2, 2% H2, 5 % H2, 20% H2, 0.5 % CH4, 1% CH4, 5% CH4, 10% CH4, and 20 % CH4 with balance helium (Douglas Fluid & Integration Technology, Prosperity, SC), which were run at each measurement timepoint to account for the PDD baseline that varied slightly each day.

### Extraction and sequencing of DNA and RNA

Biomass samples from bacterial enrichment cultures (cow, goat, digester) were collected for DNA and RNA extraction after 72 hours (Day 3) following culture transfer. DNA was collected from bacterial enrichment samples in triplicate, while RNA was collected from bacterial and fungal enrichment samples in duplicate. Hungate tube contents were centrifuged at max speed for 5 min at 4°C and biomass pellets were immediately flash frozen in liquid nitrogen. RNAlater was added to samples intended for RNA extraction prior to centrifugation. DNA was extracted from two replicates using the QIAGEN PowerSoil^®^ DNA Isolation Kit (Qiagen, Hilden, Germany) and one replicate using a CTAB protocol. This was done to improve differential coverage binning, where only duplicate DNA samples from the PowerSoil^®^ Kit extraction were used for assessing MAG relative abundance. RNA was extracted from duplicate samples using the QIAGEN RNeasy^®^ Mini Kit with on-column DNAase 1 digestion (Qiagen, Hilden, Germany). DNA and RNA sequencing was performed on the Illumina MiSeq System (v2, 600 cycles) using the Nextera XT DNA Library Preparation Kit (Illumina, CA, USA) and the NEBNext^®^ Ultra II Directional RNA Library Prep Kit at the Biological NanoStructures Lab at UC Santa Barbara. For bacterial metatranscriptomes, rRNA was depleted using the NEBNext^®^ rRNA Depletion Kit (New England Biolabs, MA, USA). For fungal metatranscriptomes, enrichment of mRNA and separation from rRNA was achieved using the NEBNext® Poly(A) mRNA Magnetic Isolation Module. Strand-specific cDNA libraries were prepared with Invitrogen SuperScript II Reverse Transcriptase (Thermo Fisher Scientific, MA, USA). Nucleic acid quantity and quality were determined using a Qubit fluorometer (Thermo Fisher Scientific, MA, USA) and an Agilent TapeStation system (Agilent, CA, USA), respectively.

### Metagenomic assembly, binning, metabolic reconstruction

Raw paired-end DNA reads were quality-trimmed and filtered, adapter-trimmed, and contaminant-filtered using BBDuk v38.75 and read quality was assessed using FASTQC v0.11.8. Trimmed reads from the same sample (cow, goat, and digester bacterial enrichments) were co-assembled using MetaSPAades v3.15.3 [11] and mapped to assembled contigs using BBMap v38.75 with minid=0.95. Resulting contigs of at least 1000 bp were binned with MaxBin2 v2.2.7 [12], MetaBAT2 v2.1512 [13], and CONCOCT v1.1.0 [14] and dereplicated using DAS-Tool v1.1.213 [15]. Metagenome-assembled genomes (MAGs) were taxonomically classified with GTDB-Tk v2.1.1 (release 9514) [16]. Bacterial MAGs were manually assigned a gram stain based on the taxonomic classification and literature references. Subsequently, gram stain specific prediction of protein product localization for each open reading frame was performed using PSORTb v3.0 [17]. Estimates of MAG completeness and contamination was performed using CheckM v1.1.2 [18].

Bacterial MAGs were annotated using the NCBI Prokaryotic Genome Annotation Pipeline version 6.10 [40]. Metabolic pathways were annotated across MAGs using gapseq v1.4.0 using the parameters “gapseq find”, “all pathways”, and “bacterial mode” [41]. Fungal reference genomes for the previously isolated and genomically sequenced *Neocallimastix lanati* [20], *Anaeromyces sp. S4* [21], *Orpinomyces sp. strain C1A* [22], *Piromyces sp. E2* [21], *Piromyces sp.finn* [21], *Neocallimastix californiae G1* [21], and *Neocallimastix frontalis var. giraffae* were retrieved from the Joint Genome Institute MycoCosm database. CAZymes and CAZyme motifs in fungal genomes and bacterial MAGs were predicted using dbcanLight v1.0.2 [23] using ‘cazyme mode’ and ‘sub mode’ options. If multiple distinct CAZyme motifs or substrate targets were predicted for the same open reading frame predictions were concatenated into a single feature.

### Metatranscriptomic analysis

Complementary DNA (cDNA) reads were quality filtered as described above for DNA. SortMeRNA [24] was used to remove rRNA sequences using multiple databases for RNA sequences. The remaining non-rRNA reads were used for mapping. RNA reads from fungal enrichment cultures were aligned to a combined fungal reference genome (concatenation of all retrieved fungal reference genomes listed above) using the bbwrap.sh command from BBMap v38.86 with the usejni parameter set to true, ten threads and all other parameters set to their default values. Bacterial RNA reads were aligned competitively against the total metagenomic assembly using the parameter minid=0.95 and then analyzed per MAG. Aligned reads were counted using the featureCounts command from Subread v2.0.3 using ten threads and all other parameters set to their default values.

For fungal genomes and bacterial MAGs, read counts from each sequencing sample were normalized for depth and gene length by calculating Transcripts Per Million (TPM) by Equation 1.

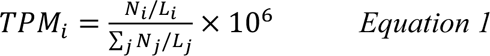

where TPMi is the TMP for transcript i, Ni is the number of reads mapped to transcript i and Li is the length of transcript i.

TPM values assigned to the same CAZyme motif, in the same genome and from the same sample (and for bacterial MAGs, the same localization) were summed. The mean TPM values across replicate samples were calculated and used for downstream analysis. For each source (digester, cow or goat), bacterial MAGs in the upper quantile of CAZyme expression, and CAZyme features predicted by PSORTb to be extracellular, were retained for differential expression analysis. Similarly, the two fungal genomes with the highest CAZyme expression from either source were retained. Differential expression analysis was performed separately on bacterial MAGs and fungal genomes from the different sources using a Wilcoxon rank sum test (wilcoxoauc function from the presto R-package, v1.0.0). Predicted CAZyme and substrate target features were mapped to the read count data, generating a CAZyme expression matrix.

## Results

### Anaerobic fungal and bacterial enrichments efficiently degrade lignocellulose to VFAs

To establish different fungal and bacterial consortia capable of degrading crude lignocellulose into VFAs, we first cultivated parallel anaerobic enrichment cultures on 1% milled sorghum in chemically defined M2 media (see Methods). Cultures were inoculated anaerobically with either cow or goat fecal pellets, or with sludge from an anaerobic digester in the presence of either chloramphenicol (selecting for fungi) or 2-bromoethanesulfonate (BES, selecting for bacteria) and passaged every 3-4 days over multiple generations (Figure 1A). In total, inoculation of sorghum media with cow and goat feces produced stable fungal and bacterial enrichments from each source, whereas inoculation with anaerobic digester sludge only produced a bacterial enrichment, suggesting the source digester had low (or no) anaerobic fungal abundance.

**Figure 1.**
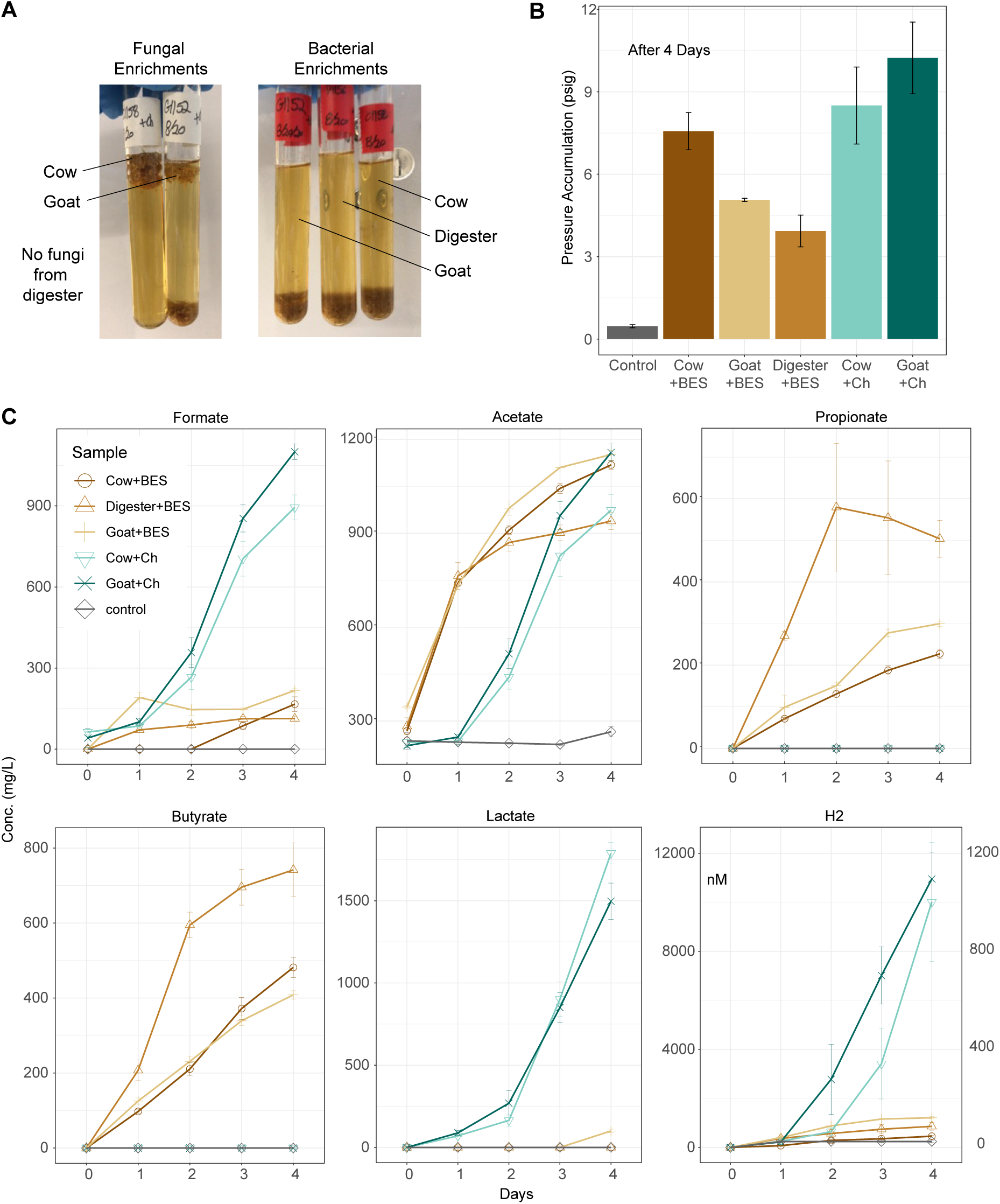
Fungal and bacterial enrichment cultures converting lignocellulosic biomass to VFAs. A) Enrichment of anaerobic fungal and bacterial consortia from the cow and goat rumen and an anaerobic digester. B) Accumulated gas production (psig) after 4 days and C) VFA, lactate, and H_2_ production by anaerobic fungal and bacterial enrichments sourced from the cow and goat rumen and an anaerobic digester. VFA and lactate concentrations reported in mg/L, H_2_ concentration reported in nanomolar. H_2_ concentrations for fungal and bacterial enrichments shown on left and right y-axes, respectively.

All enrichments were capable of sorghum deconstruction based on large increases in pressure accumulation during 4-day Hungate tube (i.e., batch) fermentations (Figure 1B). Fermentation products produced by the fungal enrichments were consistent with a bacterial-like mixed acid fermentation pathway previously reported for anaerobic fungi [25]. Both the cow and goat fungal enrichments produced high yields of formate, acetate, lactate, and hydrogen gas (Figure 1C). In comparison, the three bacterial enrichments from the cow, goat, and digester inocula produced acetate, propionate, and butyrate (C2-C4 VFAs) as main fermentation products, as well as a small amount of hydrogen gas (Figure 1C). These results are consistent with previous studies that quantified VFA production during lignocellulose degradation in anaerobic bacterial fermentations [5–7].

### Genome-centric metatranscriptomics reveals key microbes active across microbiomes

To identify key microbes responsible for lignocellulose degradation and conversion across the fungal and bacterial enrichments we recovered genomes of individual species and profiled their gene expression using metatranscriptomics.

A total of 92 MAGs were recovered across the cow, goat, and digester bacterial enrichments affiliated with diverse bacteria belonging to the phyla Actinomycetota, Bacillota, Bacteroidota, Desulfobacterota, Fusobacteriota, Pseudomonadota, and Spirochaetota, and Synergistota. A summary of the genome statistics and taxonomy of these MAGs can be found in Supplementary Table 1. Mapping of DNA and RNA reads back to these genomes revealed that MAGs affiliated with the Lachnosporaceae and Bacteroidetes dominated in abundance and activity across all bacterial enrichments (Figure 2A). In particular, several MAGs seemed to be especially important to microbiome function based on their relative DNA and RNA abundance, including Lachnosporaceae-1C (22% DNA; 30% RNA), Prevotella-1C (12%DNA; 8% RNA), and Anaeroplasma-1C (2% DNA; 7% RNA) from the cow enrichment, Ruminococcaceae-1G (3% DNA; 17% RNA), Lachnosporaceae-4G (12% DNA; 16% RNA), Lachnosporaceae-5G (3% DNA; 8% RNA), Lachnosporaceae-7G (9% DNA; 5%RNA), Bacteroidales-1G (3% DNA; 11% RNA), and Desulfovibrio-2G (1% DNA; 5% RNA) from the goat enrichment, and Bacteroides-3D (25% DNA; 27% RNA), Bacteroides-1D (25% DNA; 7% RNA), and Clostridium-1D (9% DNA, 33% RNA) from the digester enrichment (Figure 2A). Interestingly, several of the Bacteroides MAGs across the enrichments (e.g., Bacteroides-1C, Bacteroides-3G, and Bacteroides-2G) had very low relative RNA abundance compared to their relative DNA abundance (Figure 2A). This likely reflects temporal differences in activity across the 4-day batch fermentation as RNA was collected on Day 3, suggesting these species were more active during the early stages of lignocellulose breakdown.

**Figure 2.**
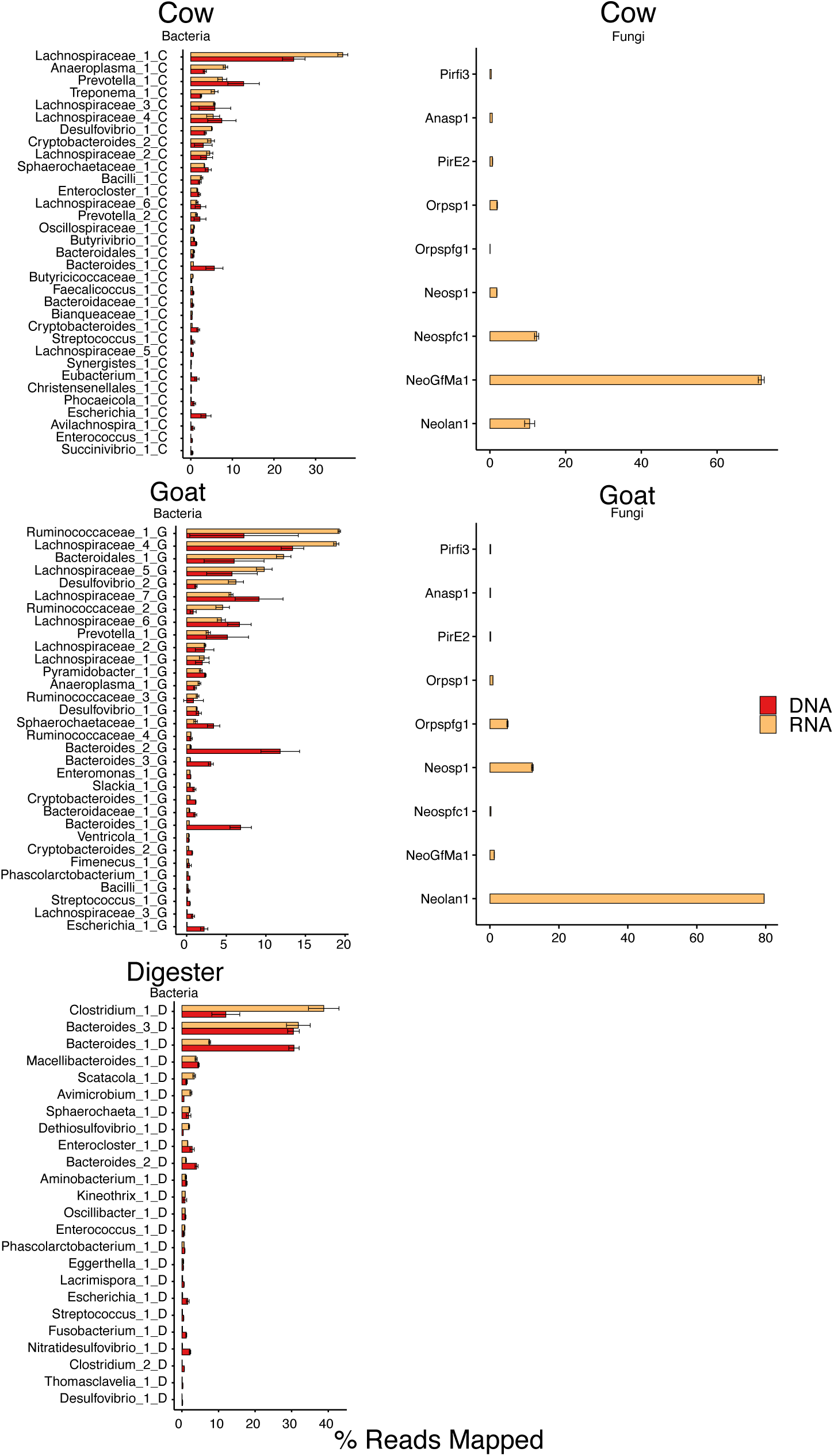
Abundance and activity of bacterial MAGs and fungal isolate genomes recovered from enrichments. Percentage of raw metagenomic (DNA - red) and metatranscriptomic (RNA - pink) reads mapped to recovered MAGs (A) and existing fungal isolate genomes (B). Fungal isolate genomes labels: Anasp1 - *Anaeromyces robustus*, Neosp - *Neocallimastix californiae G1*, NeoGfMa1 - *Neocallimastix frontalis var. giraffae*, Neolan1 - *Neocallimastix lanati*, Orpsp1 - *Orpinomyces sp*., PirE2 - Piromyces sp. E2, Pirfi3 - *Piromyces finnis*.

To identify active species from the fungal enrichments, RNA reads were mapped back to genomes available at the time from seven isolated anaerobic fungi, including *Anaeromyces robustus* (Anasp1), *Neocallimastix californiae* G1 (Neosp1), *Neocallimastix frontalis var. giraffae* (NeoGfMa1), *Neocallimastix lanati* (Neolan1), *Orpinomyces* sp. (Orpsp1), *Piromyces* sp. E2 (PirE2), and *Piromyces finnis* (Pirfi3). A large fraction of RNA reads mapped successfully to these fungal references (>95%), confirming substantial transcriptional activity from anaerobic fungi in both cow and goat enrichments (Figure 2B; Supplementary Data 1). However, due to extensive sequence conservation and gene family expansion among anaerobic fungal genomes, many reads mapped ambiguously across closely related taxa, limiting confident assignment to specific species or genomes. Consequently, downstream analyses focused on functional profiles, especially CAZy family expression, rather than species-resolved transcript abundances. At a higher taxonomic level, read mapping patterns were broadly consistent with transcriptional activity from *Neocallimastix*-dominated enrichments (Figure 2B), nevertheless high sequence similarity among genomes precluded robust species-level resolution.

### Carbohydrate active enzymes (CAZyme) expression profiles indicate substrate specialization and niche partitioning during lignocellulose degradation

Sorghum biomass is predicted to consist of crystalline and amorphous cellulose that interacts with xylan containing frequent and irregular arabinosyl substitutions. The monosaccharide composition of non-cellulosic sorghum components in mole percent is approximately 63% xylose, 17% glucose, 12% arabinose, 4% galactose, 3% galacturonic acid, and 1% glucuronic acid (Gao et al., 2020). This lignocellulosic matrix represents the primary carbon and energy substrates available to the enrichment cultures and was expected to drive microbiome structure and function [26]. To link the observed lignocellulose degradation activities to potential enzyme functions we analyzed the expression of carbohydrate active enzymes (CAZymes) across all bacterial and fungal samples. This revealed a wide range of expressed CAZymes, targeting diverse glycosidic linkages present in cellulose, hemicellulose, starch, and pectin, especially cellulase (GH9), cellulose 1,4-β-cellobiosidase (GH48/GH6), and β-glucosidase (GH3) involved in cellulose degradation and endo-1,4-β-xylanase (GH10/GH11), α-N-arabinofuranosidase (GH43), xylan 1,4-β-xylosidase (GH43), and α-L-arabinofuranosidase (GH51) involved in hemicellulose degradation (Supplementary Table 2) that have previously been implicated in lignocellulose breakdown in the cow rumen [27].

Across the rumen (cow and goat) and digester enrichments, bacterial MAGs affiliated with Clostridia (Lachnospiraceae*, Ruminococcus*) and Bacteroidia (*Bacteroides*, *Macellibacteroides*, *Prevotella*) were the main contributors to CAZyme expression (Figure 3), consistent with their high abundance and activity based on read mapping (Figure 2A). In the digester bacterial enrichment, *Clostridium*-1D was implicated as a key lignocellulose degrader with versatile hydrolytic activities based on expression of extracellular CAZymes involved in cellulose hydrolysis (GH9), xylan backbone cleavage (GH10/GH11), arabinoxylan debranching (GH43/GH51), and ester bond hydrolysis between ferulic acid and arabinose moieties in arabinoxylan (CE1) (Figure 3, Supplementary Table 3). As all these enzymes were extracellular, byproducts of degradation (xylo-oligosaccharides, sugar monomers, etc.) likely became available to other microbiome members.

**Figure 3.**
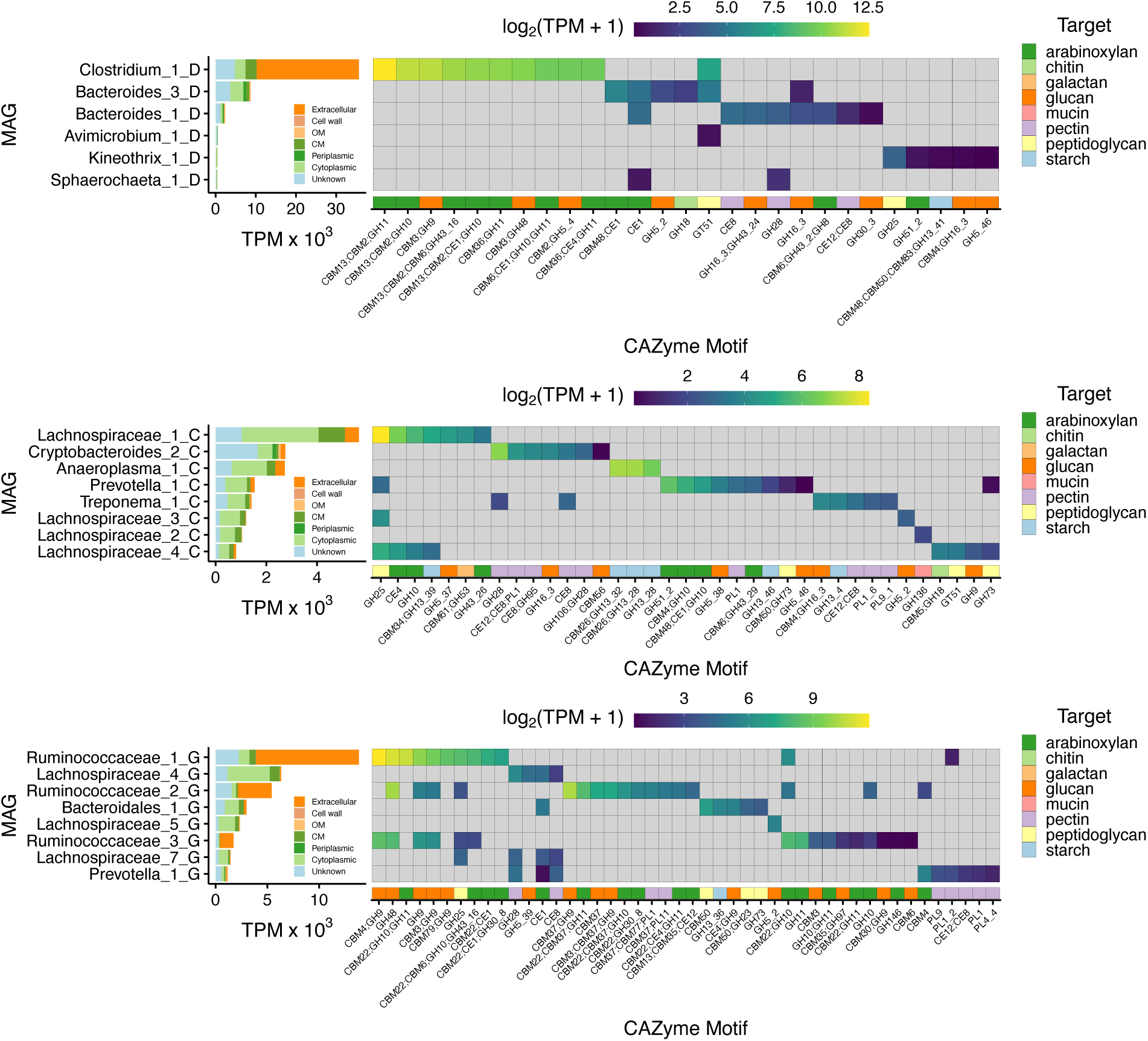
CAZyme expression across bacterial MAGs from (A) digester enrichment, (B) cow enrichment, (C) goat enrichment. Left panel – RNA expression (transcript per million, TPM) of annotated CAZymes based on predicted subcellular localization using PSORTb v3.0. Right panel – log_2_ TPM expression across CAZyme motifs per MAG. Target substrate of CAZyme motif (bottom row) predicted using dbCAN3.

In the goat bacterial enrichment, lignocellulose degradation activities appeared more distributed. Diverse *Ruminococcus* MAGs (*Ruminococcus*-1G*, Ruminococcus*-2G*, Ruminococcus*-3G) in the goat enrichment dominated CAZyme expression, including a wide array of CAZymes involved in cellulose and arabinoxylan hydrolysis (Figure 3, Supplementary Table 3). The three *Ruminococcus* MAGs expressed CAZymes with similar glycolsyl hydrolase motifs, such as GH9/GH48 (glucan) and GH10/GH11 (arabinoxylan), but differed in the carbohydrate binding domains. This suggests that in the goat enrichment, *Rumminococcus* MAGs were broadly competing for arabinoxylan but could coexist through niche specialization which arose from differing modes of access to the lignocellulosic biomass. For example, although all three *Ruminococcus* MAGs expressed cellulases from the GH9 family, *Ruminococcus*-1G and *Ruminococcus*-3G expressed GH9 linked to a CBM4, while *Ruminococcus*-2G expressed GH9 linked to a CBM37, indicating that these MAGs bind to differing sites on the glucan fraction of sorghum.

Lignocellulose degradation activities were expresedsed across an even broader set of taxa in the cow bacterial enrichment, including Lachnospiraceae*-*1C, *Prevotella-1C*, and Bacteroidales-2C. Lachnospiraceae-1C and *Prevotella*-1C appeared to be the primary degraders of arabinoxylan, while *Cryptobacteroides*-2C and *Anaeroplasma*-1C targeted pectin and starch, respectively. Lachnospiraceae-1C, which had the highest expression of total and extracellular CAZymes, was highly enriched for extracellular GH25 (lysozyme) which targets bacterial cell walls. The simultaneous expression of GH25, and preference for arabinoxylan targets, by Lachnospiraceae-1C and *Prevotella*-1C suggests that the two MAGs were broadly competing for arabinoxylan but may coexist through niche specialization conferred through diverse CBMs and debranching activities. For example, Lachnospiraceae-1C and *Prevotella*-1C both express CAZymes from the GH10 family, but GH10 CAZymes expressed by *Prevotella*-1C are further linked to binding domains (CBM4, CBM 48) and esterases (CE1), which are not associated with the GH10 CAZymes from Lachnospiraceae-1C.

Across bacterial enrichments, several MAGs displayed high expression of multifunctional enzymes consisting of several glycosyl hydrolases, glycosyl hydrolases with carbohydrate esterases, or glycosyl hydrolyses with carbohydrate binding modules (Figure 3, Supplementary Table 4). For example, *Clostridium*-1D had high expression of a CAZyme encoding two endo-β-1,4-xylanases (GH10/GH11) with a carbohydrate esterase (CE1) that may improve hydrolysis of feruloylated arabinoxylan or oligosaccharides, whereas *Ruminococcus-1G* expressed CAZymes encoding Endo-β-1,4-xylanase (GH10) with a α-L-arabinofuranosidase (GH43) as well as a glucuronoarabinoxylan endo-1,4-beta-xylanase (GH30) with a carbohydrate esterase (CE1) that also likely target branched arabinoxylans (xylan with arabinosyl and feruloyl substitutions). These multifunctional enzymes were MAG specific, especially when comparing gene expression across abundant Lachnospiraceae MAGs recovered from the different enrichments that encoded widely varying lignocellulose hydrolysis capabilities, with *Clostridium*-1D appearing to be the most robust degrader. Taken together, these results highlight different degradation strategies between bacterial enrichments sourced from diverse inocula (digester sludge, cow feces, goat feces) reflective of niche partitioning and substrate specialization.

Anaerobic fungal enrichments from the cow and goat samples also had high expression of diverse CAZymes. The repertoire of expressed CAZymes in the goat and cow enrichments were highly similar and primarily targeted arabinoxylan and glucan fractions of the lignocellulosic biomass. This suggested that, although different fungi dominated, the cow and goat fungal enrichments performed similar biochemical functions shaped by sorghum as a substrate. Compared to bacterial enrichments from the same source, the fungal enrichments consistently had a lower diversity of CAZymes, especially glycosyl hydrolases (GHs) (Figure 4). However, when only considering bacterial transcripts predicted to be extracellular, the diversity of GHs expressed by bacteria or fungi became more comparable. Comparing fungal CAZymes and the extracellular fraction of bacterial CAZymes across cow and goat sources, the fungal enrichments expressed 38 GHs not expressed by in the bacterial enrichments, including the exo-type cellobiohydrolase, GH6 (Figure 4; Supplementary Table 4). Conversely, the bacterial enrichments expressed 11 GHs not expressed by the fungal enrichments, such GH51_2, an α-L-arabinofuranosidase involved in hemicellulose degradation (Figure 4; Supplementary Table 4). This may reflect different degradation strategies between anaerobic fungi and bacteria that could be complementary, as both are well known to co-exist in the rumen.

**Figure 4.**
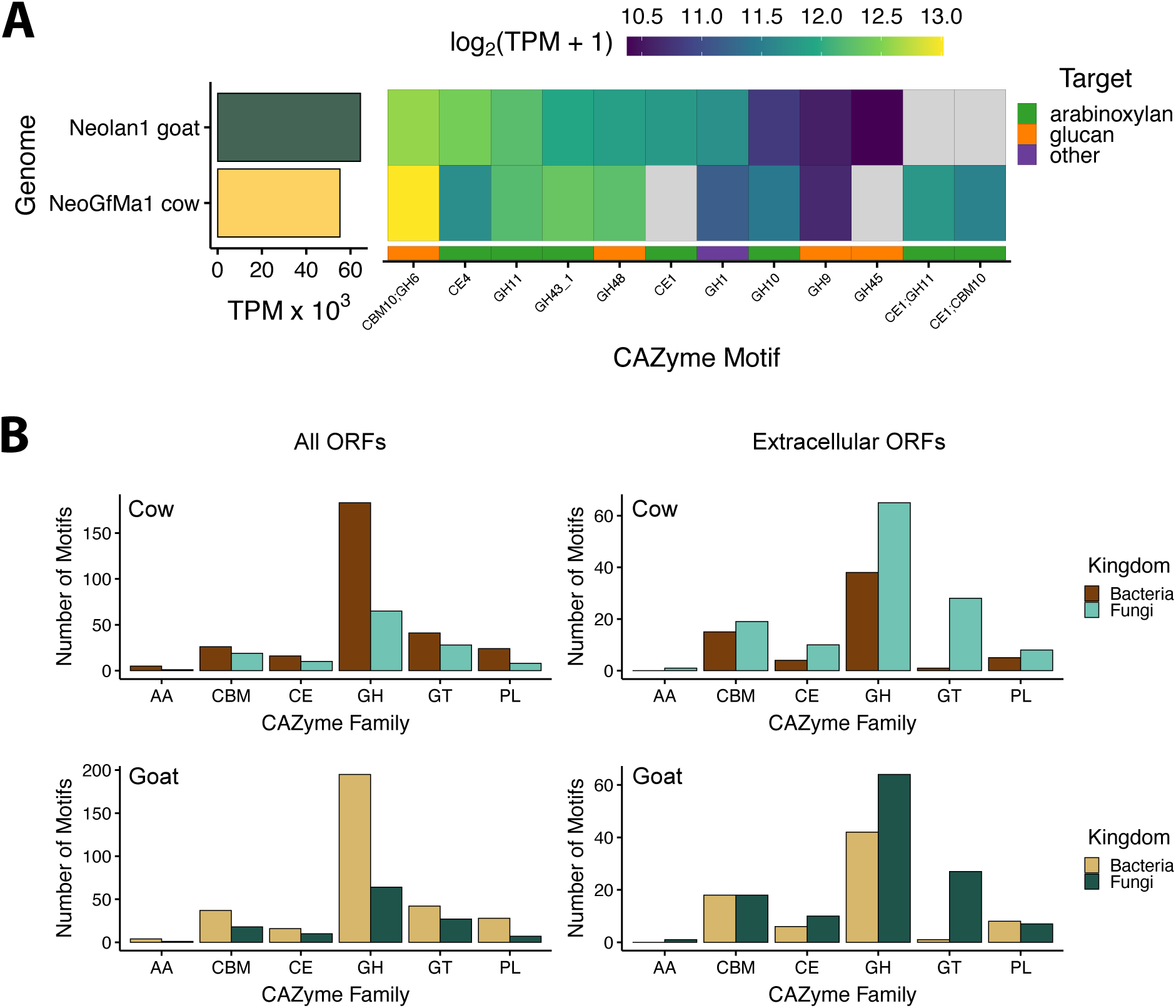
CAZyme expression across fungal reference genomes and comparison to bacterial MAGs. (A) RNA expression (log_2_ TPM) of CAZyme motifs across abundant fungal isolate genomes. Target substrate of CAZyme motif (bottom row) predicted using dbCAN3. (B) Number of motifs annotated per CAZyme family between bacterial MAGs and fungal isolate genomes. AA - Auxiliary Activities, CBM – Carbohydrate Binding Motifs, CE - Carbohydrate Esterases, GH - Glycoside Hydrolases, GT - GlycosylTransferases, PL - Polysaccharide Lyases.

Across all bacterial and fungal enrichments, genes encoding canonical lignin-depolymerizing enzymes, including lignin peroxidases, manganese peroxidases, and β-etherases, were not detected. Instead, transcripts encoding carbohydrate esterases (CE1) were detected across several abundant bacterial MAGs and anaerobic fungal genomes (Figure 3, Supplementary Data 2 and 3), consistent with the potential to hydrolyze feruloyl- and *p*-coumaroyl ester linkages associated with sorghum arabinoxylan. Canonical pathways for the subsequent catabolism of released ferulic and/or *p*-coumaric acids (e.g., feruloyl-CoA synthetase, vanillate O-demethylase, or benzoyl-CoA degradation pathways) were not detected. However, phenylacetate-CoA ligase genes were across multiple bacterial MAGs, and near-complete phenylacetate catabolic pathways were detected in a small number of low-abundance MAGs (Enterobacteriaceae-1D, Shigella-1C; Supplementary Data 4). Together, these results indicate that lignin was not extensively depolymerized under the enrichment conditions, but instead functionally circumvented through uncoupling of lignin-carbohydrate complexes. Importantly, this analysis does not preclude lignin modification via non-canonical or uncharacterized anaerobic mechanisms, as previously demonstrated by direct structural analyses of lignin during anaerobic fungal growth [42].

### Metatranscriptomic analysis reveals key fermentation pathways and microbes involved in VFA production from lignocellulosic sugars

To examine the metabolic networks of different fungal and bacterial consortia degrading lignocellulose into VFAs we reconstructed anaerobic fermentation and respiration pathways across the recovered genomes and analyzed their gene expression. MAGs were assigned to different functional guilds based on gene expression of key enzymes and metabolic pathway responsible for specific substrate utilization and expected fermentation/respiration products. This analysis revealed five major guilds across the bacterial enrichments, including hydrolytic propionate-producing bacteria, hydrolytic butyrate-producing bacteria, hydrolytic alcohol-producing bacteria, succinate/lactate producing bacteria, and sulfate-reducing bacteria.

The most abundant and active MAGs were hydrolytic bacteria involved in complex polysaccharide fermentation to propionate or butyrate, together with acetate and H_2_ formation. Propionate production was dominated by MAGs affiliated with the order Bacteroidales, including *Prevotella*-1C and *Prevotella*-2C from the cow enrichment, *Bacteroides*-2G, Bacteroidales-1G, *Bacteroides*-1G, and *Prevotella*-1G from the goat enrichment, and *Bacteroides*-3D, *Bacteroides*_1D, and *Macellibacteroides*-1D from the digester enrichment (Figure 5; Supplementary Dataset 4). These MAGs all expressed enzymes of the methylmalonyl-CoA pathway (Supplementary Dataset), including a sodium ion pumping methylmalonyl-CoA decarboxylase [28], that couples propionate formation with the generation of an ion motive force that can be used for ATP synthesis. These MAGs also had high expression of pathways for sugar utilization (glucose, xylose, and arabinose) (Figure 5; Supplementary Dataset 1) consistent with their high expression of CAZymes involved in liberating these sugars during cellulose and hemicellulose degradation (Figure 3).

**Figure 5.**
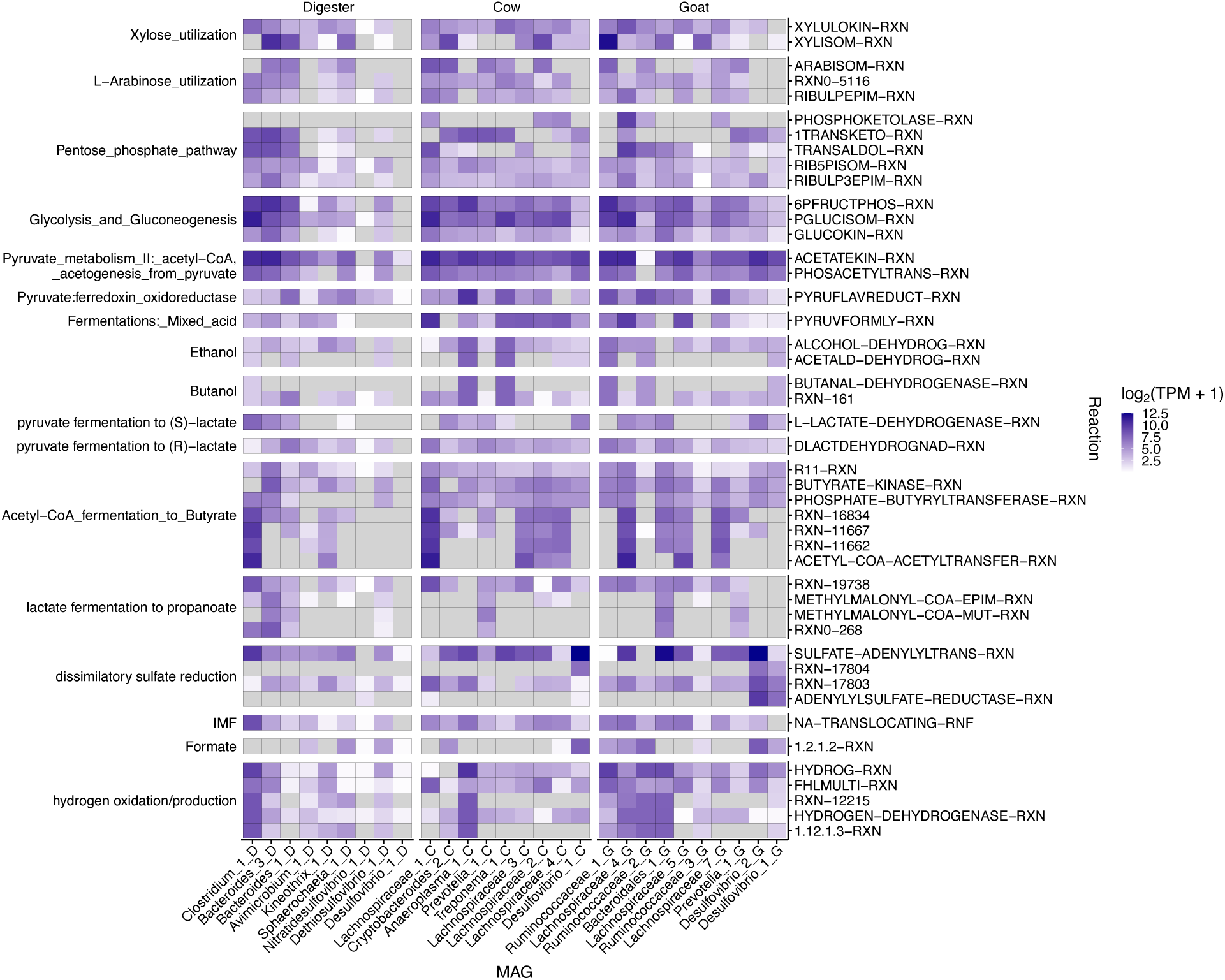
Expression of key metabolic pathways across MAGs. Rows indicate RNA expression (log_2_[TPM]) of MAG open reading frames annotated as MetaCyc reactions across key metabolic pathways via gapseq. Columns indicate bacterial MAGs from digester enrichment (left), cow enrichment (middle), and goat enrichment (right).

Butyrate production across the enrichments was dominated by MAGs affiliated with the Lachnospiraceae (Figure 5). All Lachnospiraceae MAGs had high gene expression of the reverse beta-oxidation pathway, including an electron bifurcating acyl-CoA dehydrogenase complex [29]. They also expressed the terminal enzymes phosphate butyryltransferase and butyrate kinase that couples butyrate synthesis with substrate level phosphorylation, except for *Clostridium*-1D, which instead expressed a butyryl-CoA:acetate CoA transferase [30]. While all Lachnospiraceae MAGs expressed diverse CAZymes (Figure 3) and sugar utilization pathways (Figure 5), Clostridium-1D from the digester enrichment especially stood out among the Lachnospiraceae MAGs as a specialized cellulose and hemicellulose degrader based on its high copy number and gene expression of 1,4-beta-xylanases, α-L-arabinofuranosidases, and cellulases (Figure 3).

In addition to propionate and butyrate, several MAGs affiliated with the *Ruminococcus* and Sphaerochaetaceae were predicted to be involved in complex polysaccharide fermentation to ethanol, acetate, and H_2_ formation. *Ruminococcus*-1G and *Ruminococcus*-2G had high expression of a bifunctional acetaldehyde-CoA/alcohol dehydrogenase and NAD-dependent electron-bifurcating [FeFe]-hydrogenase (HydABC), together with CAZymes and substrate utilization pathways involved in both cellulose and hemicellulose degradation (Figures 5; Supplementary Dataset 1). This metabolism is consistent with previous batch fermentation results from other *Ruminococcus* species, such as *Ruminococcus albus* isolated from the cow rumen [31, 32]. Diverse Sphaerochaetaceae MAGs, including *Treponema*-1C, Sphaerochaetaceae-1C, *Sphaerochaeta*-1D, and Sphaerochaetaceae-1C were also implicated in ethanol, acetate, formate, and H_2_ formation based on the high gene expression of alcohol dehydrogenases (iron-containing alcohol dehydrogenase and/or bifunctional acetaldehyde-CoA/alcohol dehydrogenase) and hydrogenases (iron-only hydrogenases and/or ferredoxin hydrogenases) (Figure 5; Supplementary Dataset 1). Additionally, Sphaerochaetaceae-1G also had high expression of lactate dehydrogenase and hydroxyacid dehydrogenase suggesting they may also produce lactate as an end-product.

Aside from fermentation pathways, all enrichments contained bacteria affiliated with *Desulfovibrio sp.* predicted to be capable of anaerobic sulfate respiration to hydrogen sulfide coupled to ethanol, formate, lactate, and H_2_ oxidation. In particular, *Desulfovibrio*-1C, *Desulfovibrio*-1G, *Desulfovibrio*-2G, and *Nitratidesulfovibrio*-1D had high expression of sulfate adenylyltransferase, adenylylsulfate reductase, and dissimilatory sulfite reductase involved in dissimilatory sulfate respiration together with high expression of hydrogenases, formate dehydrogenase, lactate dehydrogenase, alcohol dehydrogenase and aldehyde dehydrogenase (Figure 5; Supplementary Dataset 1), suggesting co-utilization of ethanol, formate, lactate, and H_2_ for energy generation. This is consistent with the availability of these substrates as fermentation by-products (Figure 1).

## Discussion

The use of genome-centric metatranscriptomics allowed us to analyze the metabolism and gene expression of 5 different anaerobic enrichment cultures degrading lignocellulosic biomass to VFAs. Our results reveal that across all bacterial enrichments, MAGs affiliated with the order Bacteroidales (*Bacteroides* and *Prevotella)* and family Lachnospiraceae were dominant lignocellulose degraders that fermented biomass-derived sugars to propionate and butyrate, respectively, together with acetate and H_2_ formation. Other MAGs affiliated with *Ruminococcus* from the goat bacterial enrichment, and to a lesser extent the family Sphaerochaetaceae across all bacterial enrichments, were also implicated in biomass breakdown to ethanol, lactate, formate, acetate, and H_2_ (*Ruminococcus,* Sphaerochaetaceae), consistent with the metabolism of closely related representative isolates [33, 34]. The resulting fermentation products, especially ethanol, formate, lactate, and H_2_ were predicted to be substrates for *Desulfovibrio* MAGs that performed dissimilatory sulfate reduction across all bacterial enrichments. In the fungal enrichments, lignocellulose degradation was dominated by *Neocallimastix sp.* that employed a mixed acid fermentation pathway, producing lactate, acetate, ethanol, H_2_, and CO_2_ as major end products, consistent with the metabolism of representative isolates [20]. Taken together, these results highlight that substrate (sorghum biomass) and chemical inhibitors (chloramphenicol and BES) selected for microbiomes with similar overall functional guild structure, despite containing different taxa and starting from disparate inocula.

Detailed analysis of CAZyme gene expression across the enrichments revealed common biodegradation strategies between diverse bacteria and fungi while also highlighting potential complementarities and fine-scale functional specialization. While all bacterial and fungal enrichments expressed CAZymes necessary for cellulose and hemicellulose degradation, including those involved in cellulose hydrolysis (GH9), arabinoxylan backbone cleavage (GH10/GH11), and arabinoxylan debranching (GH43/GH51/CE1), fine-scale differences between microbes based on multi-functional enzymes linking two or more CAZymes together with diverse CBMs were widespread. Most of these multi-functional enzymes localized complementary degradation functions together, such as xylan backbone cleavage with various debranching activities, likely conferring synergistic degradation effects as shown by purified enzyme studies [35]. The association of these enzymes with diverse CBMs also suggests targeted specialization towards specific lignocellulose structures (e.g., substituted backbones or branched linkages), consistent with recent observations among human gut microbiota [36]. These results implicit fine-scale substrate specialization as a major driver of microbiome assembly, niche partitioning, and species co-existence during anaerobic lignocellulose degradation.

Hydrolysis and subsequent fermentation of lignocellulosic substrates resulted in a range of fermentation acids shaped by culture conditions. Indeed, it has been shown that the spectrum of fermentation products is highly influenced by both substrate and environmental parameters, such as pH and temperature. In our study, a neutral pH of 7-7.5 and temperature of 39°C was maintained, selecting for bacterial enrichments that produced short-chain fatty acids (acetate, propionate, butyrate) and H_2_ as major fermentation products, and fungal enrichments that produced mixed acid fermentation products (lactate, acetate, formate) and H_2_. These acids appeared to be directly produced from lignocellulose-derived sugars versus indirectly via lactate as an intermediate, which can result via cross feeding between lactate-producing species (e.g., LAB, Bifidobacteria) and lactate-utilizing species. This may have resulted from having neutral versus mildly acid pH (∼5-6) conditions, the latter of which has been shown to select for lactate cross-feeding. However, general principles on what controls the distribution of fermentation products remains unclear and represents an important knowledge gap to advance anaerobic fermentation as a platform for biorefining of specific volatile fatty acids, lactate, alcohols, and other bulk chemicals [37].

While further studies are still needed to comprehensively map the functional niches driving anaerobic lignocellulose degradation to VFAs, our results provide important insights on fine-scale substrate specialization and metabolic pathways structuring microbiome assembly and function. We anticipate that these findings will help inform efforts to develop synthetic microbiomes with tailored functionality for low-cost conversion of lignocellulosic biomass to fuels/chemicals [38]. This will require isolating and assembling novel anaerobes, such as those identified here, into defined consortia to eliminate competing metabolic pathways, as well as introducing novel functionalities via metabolic engineering [39]. Such consortia would also serve to elucidate basic principles governing microbiome assembly and function, resulting in robust anaerobic fermentation platforms for sustainable biomanufacturing.

## Acknowledgements

This research was sponsored by the U.S. Department of Energy, Office of Science through grant DE-SC0022142. The work conducted at the Joint BioEnergy Institute was supported by the U.S. Department of Energy, Office of Science, Biological and Environmental Research Program, through contract DE-AC02-05CH11231 between Lawrence Berkeley National Laboratory and the U.S. Department of Energy. The United States Government retains and the publisher, by accepting the article for publication, acknowledges that the United States Government retains a nonexclusive, paid-up, irrevocable, worldwide license to publish or reproduce the published form of this manuscript, or allow others to do so, for United States Government purposes. Any subjective views or opinions that might be expressed in the paper do not necessarily represent the views of the U.S. Department of Energy or the United States Government.

## Data Availability

Raw DNA and cDNA read data and annotated metagenome assembled genomes (MAGs) can be found on the National Center for Biotechnology Information (NCBI) website under BioProject accession no. PRJNA1379681.

## SUPPLEMENTARY INFORMATION

**Supplementary Figure 1.**
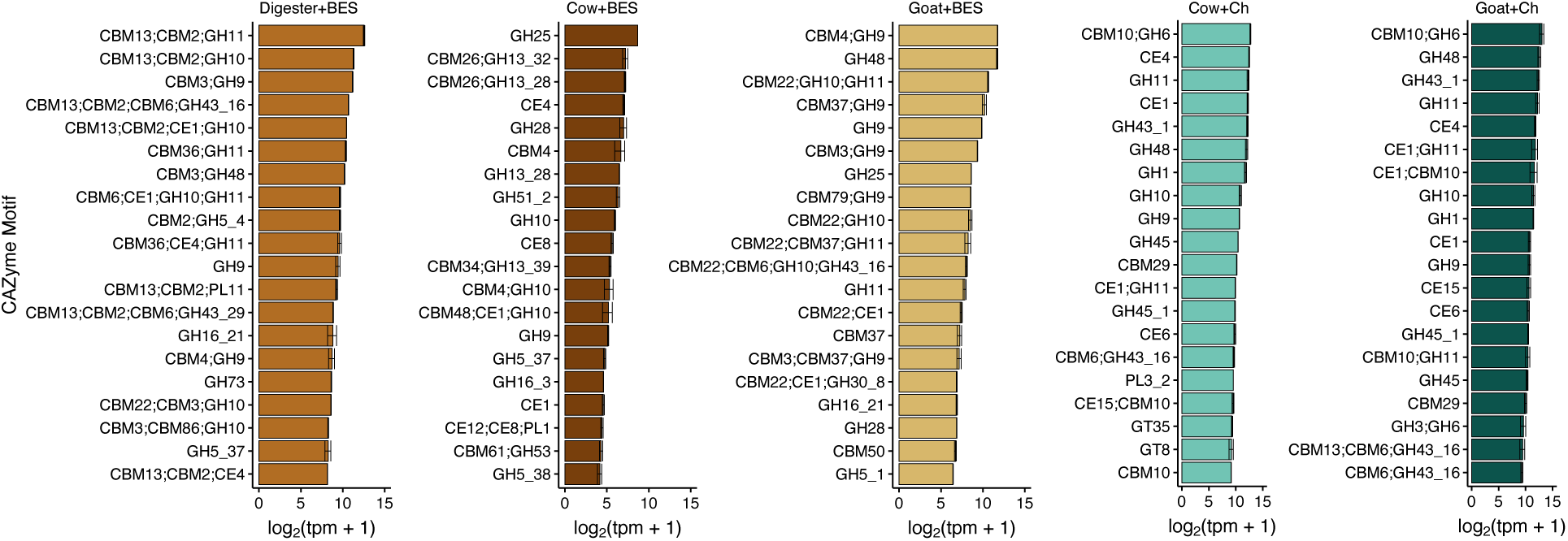
Top CAZyme motifs expressed across bacterial and fungal enrichments. CAZyme motifs were predicted using dbcanLight v1.0.2. BES – 2-bromoethansulfonate (bacterial enrichments), Ch-Chloramphenicol (fungal enrichments).

## Supplementary Datasets

**Supplementary Data 1 – MAG Statistics.** MAG statistics, including MAG size, GC content, completeness and redundancy, and DNA/mRNA read mapping, from cow (A), goat (B), and digester (C) bacterial enrichments.

**Supplementary Data 2 – Bacterial MAG CAZyme expression table.** Annotation and expression of CAZymes across all bacterial enrichment MAGs.

**Supplementary Data 3 – Fungal CAZyme expression table.** Annotation and expression of CAZymes across fungal reference genomes.

**Supplementary Data 4 – MAG pathway summary.** Predicted MetaCyc reactions and metabolic pathways across abundant MAGs recovered from cow, goat, and digester enrichments.

